# Microstructural alterations in tract development in college football: a longitudinal diffusion MRI study

**DOI:** 10.1101/2022.04.01.486632

**Authors:** Maged Goubran, Brian D. Mills, Marios Georgiadis, Mahta Karimpoor, Nicole Mouchawar, Sohrab Sami, Emily L. Dennis, Carolyn Akers, Lex A. Mitchell, Brian Boldt, David Douglas, Phil DiGiacomo, Jarrett Rosenberg, Gerald Grant, Max Wintermark, David Camarillo, Michael Zeineh

## Abstract

**Background and Objectives:** Repeated concussive and sub-concussive impacts in high-contact sports can affect the brain’s microstructure, which can be studied using diffusion MRI. Most prior imaging studies, however, employ a cross-sectional design, do not include low-contact players as controls, or use traditional diffusion tensor imaging without investigating novel tract-specific microstructural metrics.

**Methods:** We examined brain microstructure in 63 high-contact (American football) and 34 low-contact (volleyball) collegiate athletes with up to 4 years of follow-up (315 total scans) using advanced diffusion MRI, a comprehensive set of multi-compartment models, and automated fiber quantification tools. We investigated diffusion metrics along the length of tracts using nested linear mixed-effects models to ascertain the acute and chronic effects of sub-concussive and concussive impacts, as well as associations between diffusion changes with clinical, behavioral, and sports-related measures.

**Results:** Significant longitudinal increases in fractional anisotropy and axonal water fraction were detected in volleyball players, but not in football players, along with decreases in radial and mean diffusivity as well as orientation dispersion index (all findings absolute T-statistic > 3.5, p < .0001). This pattern was present in the callosum forceps minor, left superior longitudinal fasciculus, left thalamic radiation, and right cingulum hippocampus. Longitudinal group differences were more prominent and observed in a larger number of tracts in concussed (previously or in-study) football players (p < .0001), while smaller effects were noted in un-concussed players. An analysis of immediate-post concussion scans in football players demonstrated a transient localized increase in axial diffusivity, mean and radial kurtosis in the left uncinate and right cingulum hippocampus (p < .0001). Finally, football players with high position-based sub-concussive risk demonstrated increased orientation dispersion index over time (p < .0001).

**Discussion:** The observed longitudinal changes in our volleyball cohort likely reflect normal development in this age range, while the relative attenuation of these effects seen in football, and especially concussed athletes, could possibly reveal diminished myelination, altered axonal calibers, or depressed pruning processes leading to a static, non-decreasing axonal dispersion. This prospective longitudinal study demonstrates significantly divergent tract-specific trajectories of brain microstructure, possibly reflecting a concussive and/or repeated sub-concussive impact-related alteration of normal white matter development in football athletes.

## 1. Introduction

Repeated concussive and sub-concussive high-velocity or acceleration impacts may affect the brains of players in high-contact sports, i.e. American football, with possible long-term cognitive sequelae ^1–3^. Given the millions of people in the US and worldwide engaged in contact sports, it is imperative to understand the nature and time course of these brain changes. Diffusion MRI (dMRI) is an imaging modality sensitive to microstructural alterations^4^ and can noninvasively quantify subtle changes in brain microstructure along fiber tracts associated with sport impacts^5–7^.

Significant inter-subject variability in brain microstructural anatomy complicates interpretation of cross-sectional studies of high-contact players vs. controls. Identification of changes is most specific when in reference to an individual subject’s baseline imaging, which requires longitudinal rather than cross-sectional study design. However, relatively few longitudinal studies have investigated these changes in sports, even fewer incorporate baseline imaging, while studies greater than 1 year in duration are even more scarce^8,9^. This is presumably due to their longer duration and logistical complexity^10^.

The brain’s microstructural anatomy is dynamic and matures over the lifespan^9^. Furthermore, it remains unknown which is a greater contributor to potential long-term brain injury: concussion, cumulative high-velocity sub-concussive impacts, or both^11^. Dissecting apart the evolution of these changes from normal development requires not only a mix of concussive and non-concussive high-contact players in a longitudinal study, but also the inclusion of non-concussive low-contact players as a control group -which has been used sparsely in prior work.

dMRI provides a wealth of information, but at the same time is a technique with myriad challenges and many possible microstructural interpretations^12^. Even the most commonly studied metric, fractional anisotropy, can have multiple interpretations as a basis for the changes^13,14^. While the diffusion tensor imaging (DTI) model is the most commonly used in clinical and research settings, advanced signal representation or multi-compartment modeling techniques, such as diffusion kurtosis imaging (DKI), neurite orientation dispersion and density imaging (NODDI)^15^, and white matter tract integrity (WMTI), attempt to distinguish intra- and extra-cellular or linear from non-linear components of diffusion signals. These more sophisticated modeling techniques have the potential to disentangle such contradicting changes and better discern the underlying microstructural phenomena^4^.

Concussive and sub-concussive impacts may differentially affect specific brain tracts or portions of tracts. Furthermore, these effects may interact with developmental and longitudinal neural processes, such as myelination and pruning^16^. Thus, imaging studies need to pinpoint and investigate the location, magnitude, and potential mechanisms of (chronic and acute) tract-based microstructural alterations. However, the majority of prior studies average diffusion changes over the entire length of fiber tracts, potentially obscuring local changes, or employ voxel-wise statistics or skeletons that do not ensure voxel correspondence within the same tract across subjects^10,17,18^.

In this work, we study longitudinal alterations in brain microstructure in a high-contact college cohort (football), in comparison to a low-contact college cohort (volleyball) with baseline imaging and follow-up extending out to 4 years. We employ a two-shell dMRI acquisition, and, in addition to traditional diffusion tensor imaging (DTI) measures, analyze advanced signal representation (DKI) or multi-compartment modeling (NODDI and WMTI) metrics along the fiber tracts. We further investigate the acute and chronic effects of both sub-concussive as well as concussive impacts on tract-specific diffusion metrics. Finally, we study the associations between tract-specific diffusion changes with clinical, behavioral, and demographic measures including the Standardized Concussion Assessment Tool (SCAT) score, history of concussion, years of prior football experience and player position-based concussion risk.

## 2. Methods

### 2.1. Study Population

We prospectively enrolled 63 high-contact (football) and 34 low-contact (volleyball) college players (**Suppl. Fig. 1**) with written informed consent. All procedures were in accordance with the Health Insurance Portability and Accountability Act and Institutional Review Board. Exclusion criteria included self-reported history of brain surgery, severe brain injury, or major neurologic, psychiatric, or substance abuse disorder. In conjunction with athletic trainers, the study was introduced to team members, and all interested eligible players were enrolled. Players underwent brain MRI i) annually at the start of each season, ii) within 24-96 hours after a concussion (when possible), and iii) after the last season of sport participation. For the first year of the study, players had an additional end-of-season imaging session.

Football players self-reported their history of prior concussions (number and timing), their history of football involvement prior to enrollment in the study (number of years), and the current position played. One football athlete did not provide years of tackle football experience. Players underwent a pre-first-season SCAT (versions 2 and 3) evaluation^19^. The SCAT evaluation included the cognitive assessment, coordination, balance, and standardized assessment of concussion portions of the SCAT 2/3 (maximum score: 61).

Each athlete had 1 to 8 MRI time-points spanning a maximum of four playing years. Incidences of concussion during the study period were identified by the athletic trainers. In total, 18 concussions occurred in 15 football players, three of whom had at least two concussions. Of those concussions, none involved loss of consciousness and the mean return-to-play time was 10.3 days (excluding one athlete who did not return to play). Post-concussion MRI was captured for 13 of these 18 concussions, among 12 football players. One of these 12 had two concussions with post-concussion MRI, and only the latter scan, which had a longer return-to-play time, was used in the concussion analysis. One subject was excluded from post-concussion analysis because their MRI was done two weeks after their concussion. In one subject with postconcussion MRI, their first imaging time-point was two days post-concussion without prior baseline imaging. Four volleyball players suffered in-study concussions, and their subsequent scans were excluded from further analysis. A board and CAQ-certified neuroradiologist (MZ) blindly examined all scans for incidental abnormalities, resulting in exclusion of three players (9 scans in total).

Given the longitudinal nature of this analysis, we only included subjects with two or more scans, at least 6 months apart. In total, this resulted in the inclusion 271 scans from 73 subjects: football: 49 players – 193 scans, volleyball: 24 players – 78 scans (**Suppl. Fig. 1**).

### 2.2. Image Acquisition

A 3T MRI (GE MR 750, Milwaukee, WI, USA) protocol was acquired (**Supplementary Material**). Briefly, for the described analyses the following sequences were acquired: 1) A 1mm isotropic T1-weighted scan; 2) Multi-shell whole-brain axial 2D pulsed gradient spin echo diffusion MRI with 60/30 directions at b=2500/800s/mm^2^ and voxel size=1.875×1.875×2.0mm^3^; and 3) A 1mm isotropic gradient echo field-map for distortion correction.

### 2.3. Diffusion pre-processing

Using FSL 5.0.12, we corrected for eddy currents and motion using *eddy_cuda*, utilizing default parameters and the following options: intra-volume (slice to volume) motion correction with 5-iterations and temporal regularization of 1, quadratic spatial model of the field generated by currents, detection and replacement of outlier slices, and slice outlier definition of 3.5 standard deviations. We corrected for distortions using the de-spiked and de-noised field map using FSL’s FUGUE. All scans were visually checked for artifacts and alignment between shells (b2500/b800), and in cases where misalignment was observed an alignment step with nearest-neighbor interpolation was performed.

### 2.4. Diffusion metrics

Weighted-least squares diffusion tensor fitting yielded fractional anisotropy (FA), mean, axial and radial diffusivity (MD, AD, RD) using FSL’s *dtifit*. Neurite Orientation Dispersion and Density Imaging (NODDI)^15^ derived metrics (orientation dispersion -ODI, intra-cellular volume fraction - FICVF, free water volume fraction -FISO) were derived using the NODDI toolbox (with default parameters and sigma parameter=2.0). Kurtosis and white matter tract integrity (WMTI) metrics were calculated using the DESIGNER software^20^, which provided mean, axial, and radial kurtosis (MK, AK, RK) as well as axonal water fraction (AWF) (**Fig. 1**).

**Figure 1.**
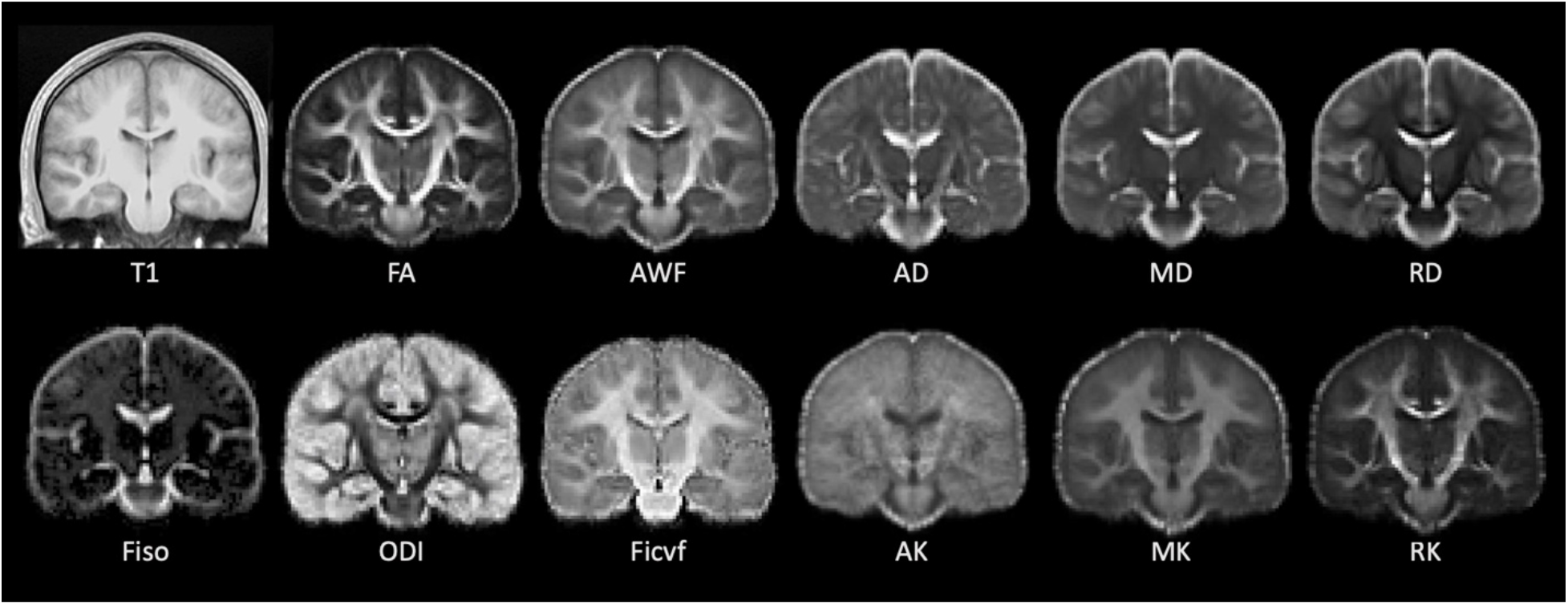
Representative images of all diffusion-based imaging metrics used in the study. Average maps of first season volleyball players in T1 space through generation of a study-specific atlas (top left). Metrics include: FA=fractional anisotropy, AWF=axonal water fraction, AD=axial Diffusivity, MD=mean diffusivity, RD=radial diffusivity, FISO=free water (isotropic) volume fraction, ODI=orientation dispersion index, FICVF=intracellular volume fraction, AK=axial kurtosis, MK=mean kurtosis, RK=radial kurtosis.

### 2.5. Whole-brain tractography

Fiber tracts in each athlete’s brain were generated using MRtrix3. *dwi2response* was used to estimate multi-shell, single-tissue response functions of the diffusion signal for each brain. *dwi2fod* was used to find fiber orientation distributions for each voxel with constrained spherical deconvolution. Probabilistic whole-brain tractography was performed using *tckgen’s iFOD2* algorithm^21^, with default parameters.

### 2.6. Automated fiber quantification (AFQ)

To quantify tissue properties along the major fiber tracts in each athlete’s brain, we applied the automated fiber quantification (AFQ) method^17^ (**Supplementary Material**). Briefly, AFQ samples diffusion metrics at equally spaced nodes along 20 major fiber tracts. This enables comparisons of different imaging metrics between individuals or groups along the extent of fiber pathways. In this work, the nodes along each tract were linearly resampled to ten nodes.

### 2.7. Statistics

A linear mixed-effects model examined changes based on fixed effects of sport (football versus volleyball), age at time of baseline scanning, years after baseline scan (time), and the interaction between time and sport. Random effects included subject ID and node number along tract (10 nodes per tract). To control for type 1 errors, we adjusted p-values across all tracts and diffusion metrics using ‘hommel’ family wise error correction using R’s *p.adjust* package (v1.6), a more conservative correction method than the typically used false discovery rate (FDR). We focused on 9 metrics previously shown to be sensitive to changes after repeated sub-concussive events^15,22,23^: DTI metrics (FA, MD, RD), NODDI metrics (ODI, FICVF, FISO), and kurtosis/WMTI metrics (MK, RK, AWF). Statistics were computed using R version 3.6.0 and additionally verified using Stata (v15.0; StataCorp LP, College Station, TX). Normality of residuals was confirmed by plotting histograms and Q-Q plots.

### 2.8. Football vs. Volleyball Analysis

Applying the mixed-effects model to the diffusion metrics, we examined: 1) baseline shifts between sports (effect of sport irrespective of time) and 2) differences in the temporal trajectories between sports (the interaction effect between sport and time). To visualize node effects (e.g. **Fig. 2**b), a three-way interaction model (sport, time, and node) was used. In this analysis, we excluded all immediate post-concussion timepoints to look at the chronic effects of injury.

**Figure 2.**
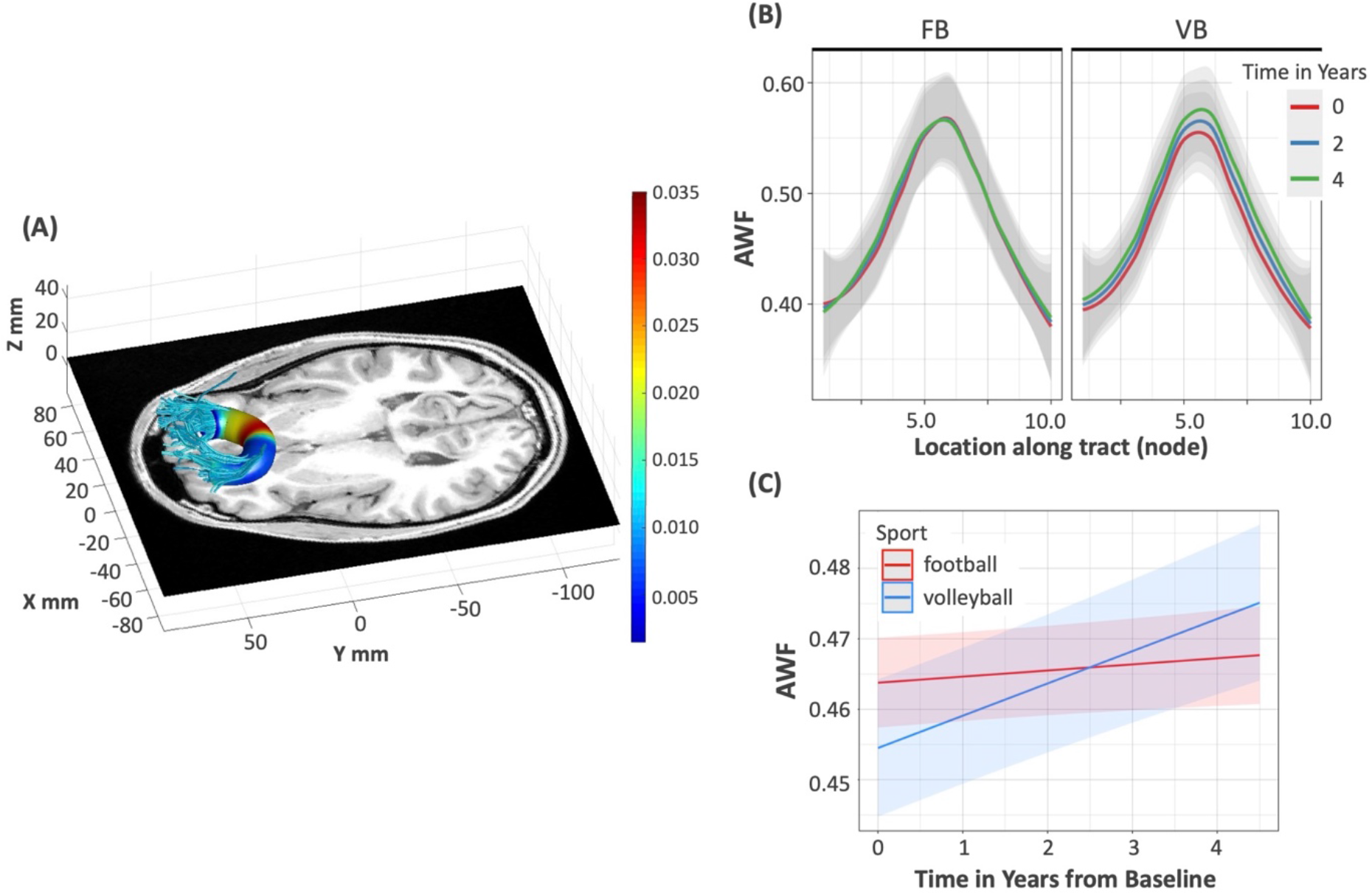
Divergent longitudinal AWF trajectory between football and volleyball players in the callosum forceps minor. (highlighted from Table 2). A) Rendering of localized effects of diverging AWF trajectories along callosum forceps minor. Tractography was performed on a representative baseline volleyball scan and overlayed on the subjects T1 image. Color coding indicates the node-wise predicted AWF difference between sports using a three way interaction model – red indicate larger, blue indicate smaller group differences/effects. B) Plot of AWF values along the tract nodes (parameterized location 1-10) for football (FB) and volleyball (VB) players, for different time points of the study (0 years – red, 2 years – blue, 4 years – green). Note the larger time-related differences in AWF VB players, especially towards the center of the tract (nodes 5-6), compared to the small divergence across the tract in FB players. C) The trajectories of AWF values over time across the whole tract, as estimated by the nested linear mixed-effects model, for football (red) and volleyball (blue) players. Note the relatively flat trajectory in football, compared to the increasing values in volleyball, the latter attributed to normal brain development, without effect of exposure to high-contact sports.

### 2.9. Concussion analyses

i. To assess chronic effects, we excluded immediate post-concussive scans, and used the above model, but separated our cohort into three groups: football players with prior or in-study concussion (N=24), football players with no prior or in-study concussion (N=25), and volleyball players (N=24). We performed all 2-way comparisons of these three cohorts.
ii. To examine acute effects of concussion in white matter microstructure, we compared immediate post-concussive scans to each subject’s most recent non-post concussive scan. This included two football athletes who were not part of the 49 because their only available MRI follow-up was less than six months apart. For one subject with post-concussion MRI, no baseline scan was available, so their follow-up scan (2.5 months later) was used as baseline. We included two additional metrics sensitive to post-concussive changes along the axons^24,25^, AD and AK.

### 2.10. Within football analyses

Examining only football athletes, we utilized a mixed-effects model to assess the following:

#### Presence of concussion

A binary variable reflected whether players had experienced either an in-study concussion and/or a concussion prior to study enrollment. This splits the football group to 25 players (100 scans) with no prior or in-study concussion, and 24 subjects (80 scans) with a prior or in-study concussion.

#### Prior football exposure

We tested whether prior experience playing tackle football before their baseline MRI was related to microstructural changes.

#### Player position

In order to test whether impact risk based on player position was related to microstructural changes, we used literature values of HITsp for each player position as a measure of impact severity^26^. HITsp is a composite measure that incorporates linear and rotational acceleration, impact duration, and impact location for each position.

#### SCAT

We tested whether microstructural changes were related to the players’ SCAT score^19^.

For each of these metrics (presence of concussion, prior football exposure, player position, SCAT score), we utilized a mixed-effects model to test if they contributed to baseline or longitudinal microstructural changes. Results were corrected using ‘hommel’ adjustment separately for each behavior but across all microstructural metrics and nodes, p<0.05, adjusted.

## 3. Results

### 3.1. Cohort Demographics

There was a small but statistically significant difference in age at baseline between football and volleyball players (football: 19.11±1.57 years, volleyball: 19.57±0.91 years, p<0.001; **Table 1**). There was a significant difference in body mass index (BMI) between football and volleyball players (football: 29.72±3.85, volleyball: 23.45±2.08, p<0.001). A subset of volleyball players had prior tackle football experience; however, it was substantially shorter compared to football players: 0.08 years (range: 0-1) vs. 8.76 years (range: 4-14) (p<0.001). Sixteen out of 49 football and two out of 24 volleyball players had concussions prior to the study. SCAT was only collected in football players, and there were no statistically significant differences in baseline SCAT scores between players with or without prior concussion. Football had very low instances of ADHD (2 players) and learning disabilities (1 player), while volleyball had none. In football there were significantly more African American players (19/49) compared to volleyball (0/24) (p<0.001), and significantly fewer Caucasian players (21/49 vs. 19/24 in volleyball) (p=0.005).

**Table 1.** Cohort Demographics. Standard deviations, as well as minimum and maximum scores, are shown for mean age at baseline, body mass index, years of prior football experience, and SCAT scores. Number of players with prior concussion, incidence of ADHD, learning disabilities, and race are detailed. SCAT scores were measured and compared within football only (players without prior concussion (left column) vs. with prior concussion (right column)). A Wilcoxon rank sum test was performed for age at baseline, body mass index, years of prior football experience, number of players with prior concussions, and SCAT scores, while a Fisher’s exact test was utilized for all other categories.

|  | Football<br>mean (standard deviation)<br>[min, max] or<br>ratio (percent) |  | Volleyball<br>mean (standard deviation)<br>[min, max] or<br>ratio (percent) | P-value |
| --- | --- | --- | --- | --- |
| Age at Baseline | 19.11 (1.57)<br>[17.61, 27.18] |  | 19.57 (0.907)<br>[18.32, 21.31] | <0.001 |
| Body Mass Index | 29.72 (3.85)<br>[21.70, 38.18] |  | 23.45 (2.08)<br>[11.27, 28.16] | <0.001 |
| Years of Prior Tackle Football Experience | 8.76 (2.57)<br>[4.0, 14.0] |  | 0.08 (0.27)<br>[0.0, 1.0] | <0.001 |
| SCAT score | Without<br>concussion | With prior<br>concussion |  | 0.472 |
|  | 54.82 (3.94)<br>[42.0, 61.0] | 55.43 (2.98)<br>[51.0, 61.0] |  |  |
| Athletes with prior concussion | 16/49 (33%) |  | 2/24 (8%) | 0.041 |
| Athletes with ADHD | 2/49 (4%) |  | 0/24 (0%) | 1.00 |
| Athletes with Learning Disabilities | 1/49 (2%)<br>*subject had dyslexia* |  | 0/24 (0%) | 1.00 |
| Caucasian | 21/49 (43%) |  | 19/24 (79%) | 0.005 |
| African American | 19/49 (39%) |  | 0/24 (0%) | <0.001 |
| Other/Mixed/Unknown | 9/49 (18%) |  | 5/24 (21%) | 1.00 |

ADHD: Attention deficit hyperactivity disorder.

### 3.2. Longitudinal differences in diffusion metrics between sports

Immediate post-concussion scans were excluded in this analysis to focus on chronic changes. Football players were found to have significantly divergent age-related trajectories in multiple microstructural metrics and tracts, compared to volleyball controls (**Table 2**; **Suppl. Table 1**). More specifically, metrics in four tracts, forceps minor of corpus callosum, left superior longitudinal fasciculus-SLF, left thalamic radiation and right cingulum hippocampus, had divergent time trajectories between football and volleyball. The temporal patterns were uniform across tracts: AWF and FA increased and MD/RD/ODI decreased in volleyball relative to football.

**Table 2.**
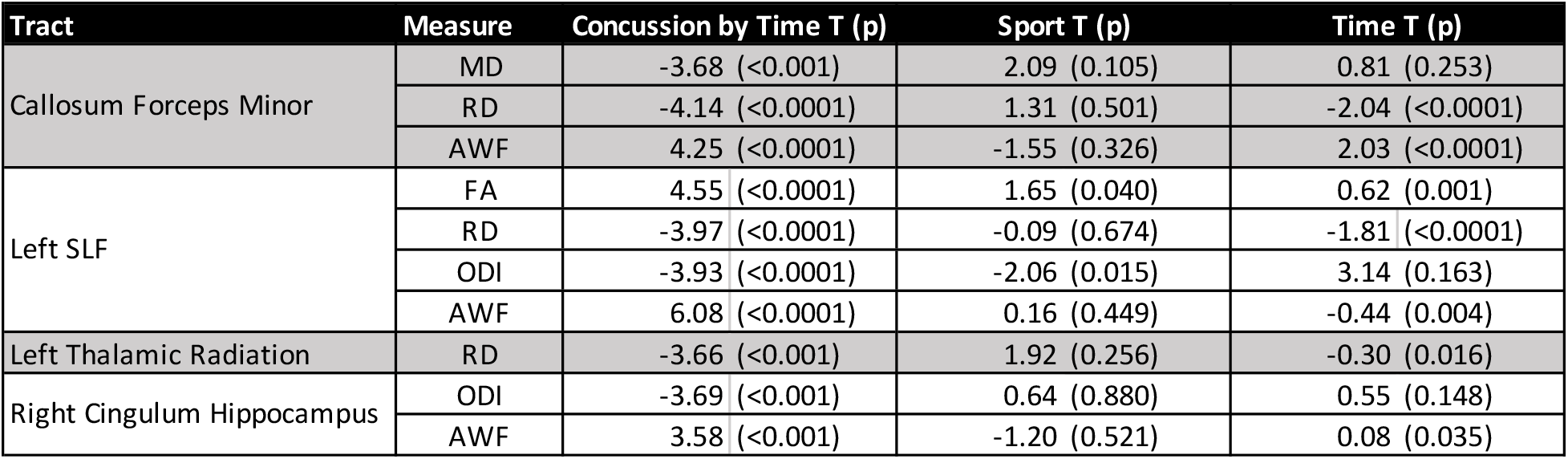
Comparison of tract-specific diffusion metrics in football vs. volleyball over time. Only metrics with significant longitudinal differences (“Sport * Time” interaction term, after FDR correction) are reported. T-statistic (p-value) for Sport, Time, and their interaction from the longitudinal linear mixed-effects models.

As an example, in the callosum forceps minor, volleyball controls showed a decreasing MD and RD and increasing in AWF over time, an effect substantially diminished in football players (**Table 2**, sport by time). **Figure 2** highlights the longitudinal AWF differences between sports in the callosum forceps minor, both within locations (nodes) along the tract (**Fig. 2**b), and in the longitudinal trajectory, based on the nested linear mixed-effects model (incorporating all the tract nodes) (**Fig. 2**c).

The left superior longitudinal fasciculus (SLF) exhibited the largest “sport-by-time” effects (for 4 diffusion metrics) (**Table 2, Fig. 3**). As seen in the nested model over time (**Fig. 3**-left), as well as in the distribution across the tract nodes (**Fig. 3**-right), the low-contact (volleyball) players showed an increased FA and AWF and decreased RD and ODI with time, but this longitudinal change was highly attenuated in high-contact (football) players. Similar findings were present in the left thalamic radiation, the right cingulum hippocampus, and the callosum forceps minor (**Table 2**). The results were unchanged when handedness was accounted for.

**Figure 3.**
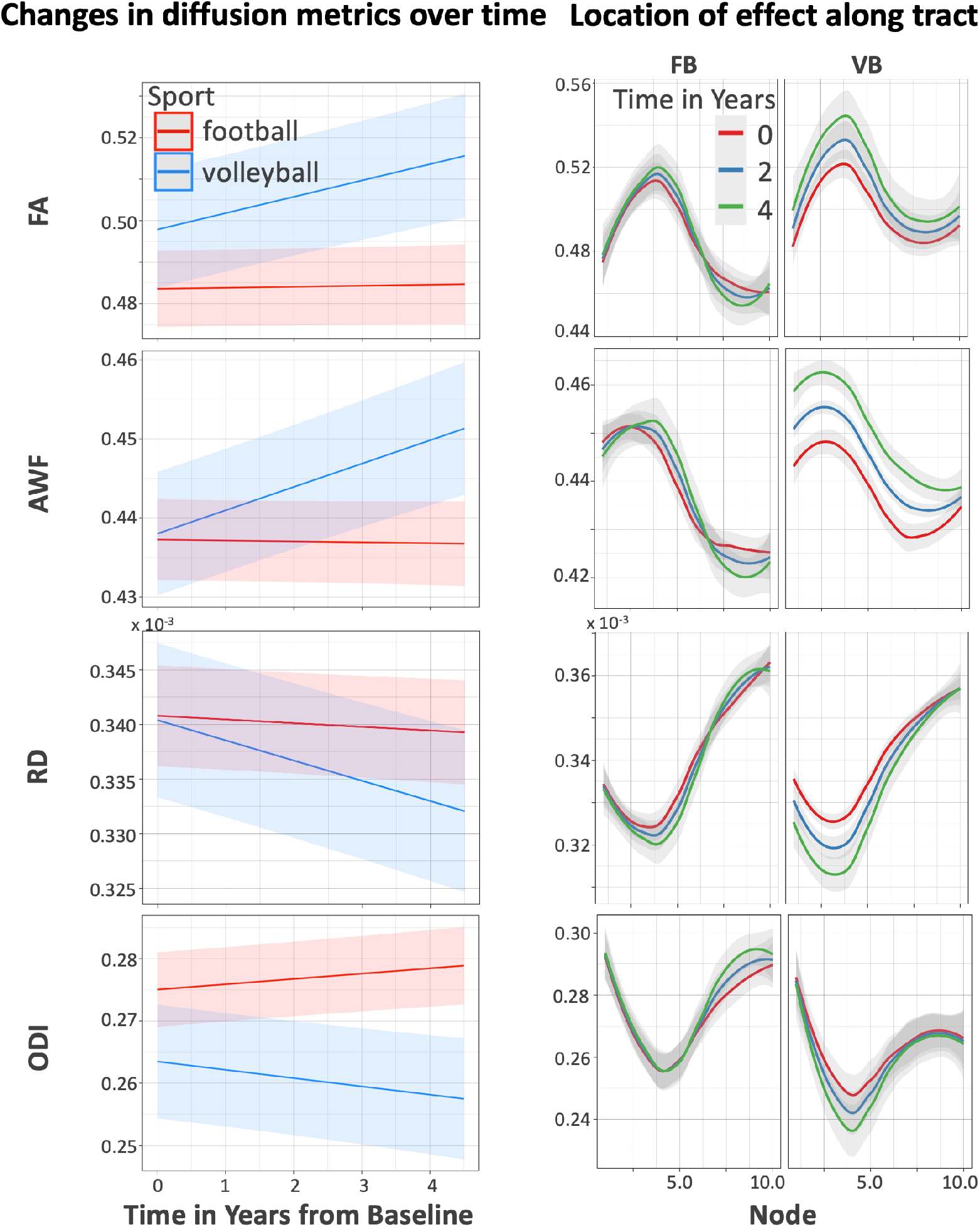
Longitudinal diffusion metrics in left superior longitudinal fasciculus (SLF), in football (FB) vs. volleyball (VB) players. Left: The trajectories of diffusion metrics values over time across the whole SLF, as estimated by the nested linear mixed-effects model, for football (red) and volleyball (blue) players. Right: The same diffusion metrics for football (FB, left column) and volleyball (VB, right column), plotted along the tract nodes (parameterized location 1-10), for 3 distinct timepoints over the course of the study (0 years – red, 2 years – blue, 4 years – green).

### 3.3. Effect of concussions

#### 3.3.1. Chronic effects of prior or in-study concussions

We sought to determine whether any prior or in-study concussion had chronic effects on tract microstructure across sports. This analysis excluded immediate post-concussive scans, analyzed in the next section. Of 49 football players, 25 did not report any concussions, 11 had prior concussions, 8 had in-study concussions and 5 had both.

Results of the nested linear mixed-effects models for the volleyball vs. concussive football comparison are summarized graphically in **Figure 4**a, with model statistics described in **Suppl. Table 2**. We observed that football players with prior or in-study concussion exhibited differences in multiple diffusion metrics and tracts when compared to volleyball controls (**Figure 4**b; **Suppl. Table 2**, “Concussive Football”). These effects were in the same direction as the ones found in the previous section (where all football were compared to volleyball players), with the same relative absence of FA/AWF increases and RD/ODI decreases over time in previously concussed football players compared to volleyball. It is worth noting that most (but not all) sport-by-time results found in the previous section are included in these results, pointing to (prior or in-study) concussions being a strong determinant of chronic microstructural alterations in the brains of high-contact sports players.

**Figure 4.**
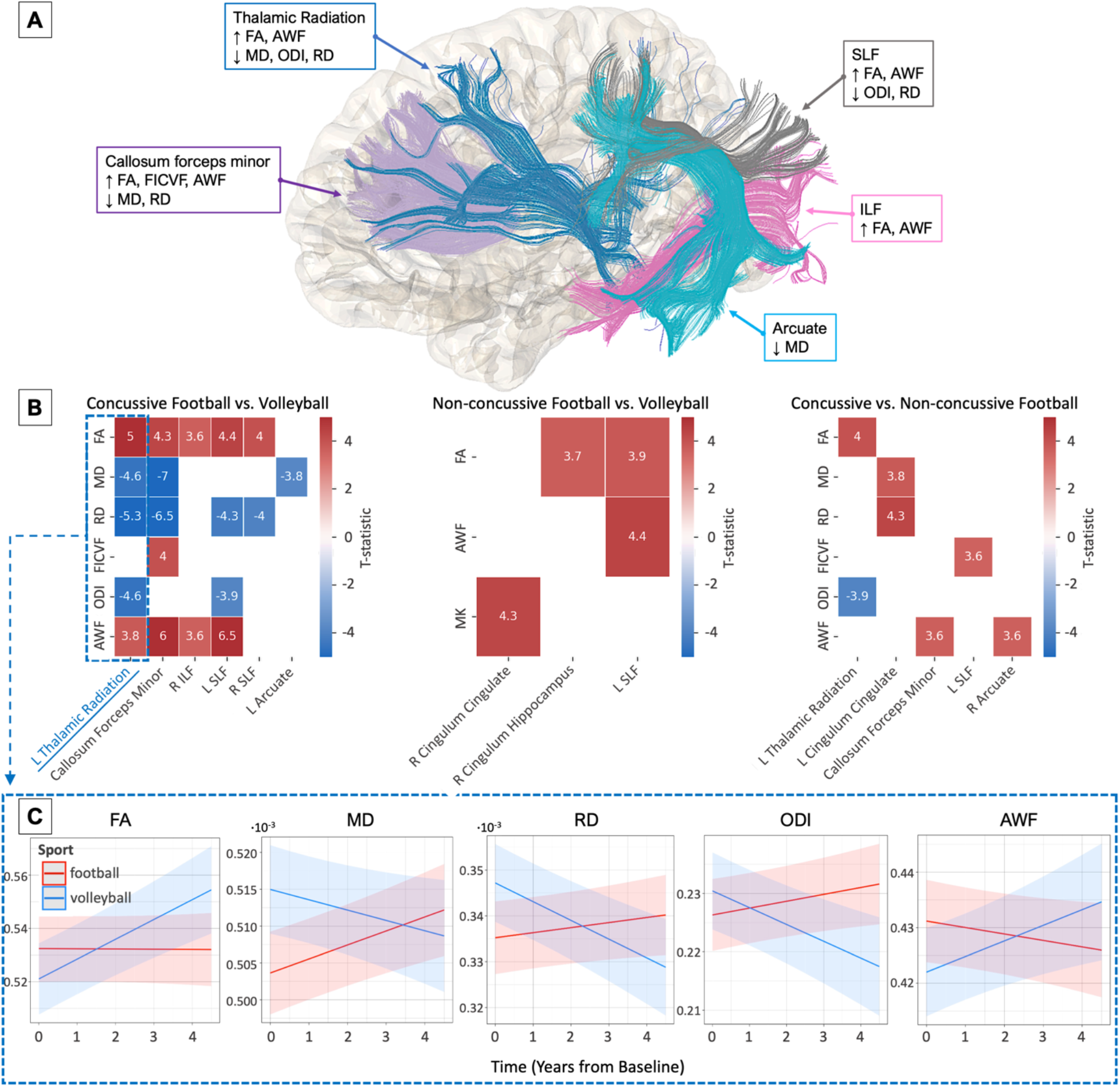
Longitudinal comparison between i) volleyball players, ii) football players with prior or in-study concussion (concussive) or iii) without concussion (non-concussive). A) Summary of tract-specific findings for volleyball vs. concussive football players. 3D rendering (using diffusion tractography) of the fiber tracts with observed longitudinal group. Each (color-coded) box includes the summary of metric changes between the two groups observed within respective tracts (shown with an arrow). Upward and downward arrows within each box represent significant longitudinal metric increases and decreases, respectively, in volleyball compared to concussive football players. For instance, in the left arcuate fasciculus (depicted in cyan), volleyball players had a significant MD decrease overtime compared to concussive football. B) Heatmaps summarizing tract-specific findings across all comparisons between the 3 groups; highlighting that longitudinal changes were more prominent and in more tracts when volleyball were compared to concussed football players (left). Heatmap values represent the T-statistic of the “Group x Time” variable from the nested mixed effect models (detailed model results summarized in **Suppl. Table 2**). Tracts are on the x-axes and diffusion metrics on the y-axes for each heatmap/group comparison. Non-significant findings are depicted as blank cells. C) Trajectories of diffusion metrics values over time across the left thalamic radiation, as estimated by the nested linear mixed-effects model, for concussive football (red) and volleyball (blue) players.

In contradistinction, football players without a history of concussion showed few differences over time against volleyball (**Figure 4**b; **Suppl. Table 2**, “Non-concussive football”). Nonetheless, longitudinal alterations were observed in the right cingulum hippocampus, not present in the “Concussive Football” analysis; suggesting that some microstructural alterations may be attributed to repetitive sub-concussive impacts only. Similarly, some changes, such as FA and AWF in the left SLF, were observed in both analyses (albeit stronger in the “With concussion” analysis); pointing towards a potential synergistic effect of concussive and repetitive sub-concussive impacts.

A direct longitudinal comparison of concussive vs. non-concussive football players (**Figure 4**b, **Suppl. Table 2**) showed that non-concussed football players have relative increases in FA/AWF and decreases RD/ODI over time compared to concussed football players. These results mirror the findings observed in the football vs. volleyball analysis, albeit with smaller effect sizes, and within fewer longitudinal group differences in metrics and tracts. Our findings support the above hypothesis, that concussions may alter the normal brain development and may be driving some of the divergent trajectories between low- and high-contact athletes.

#### 3.3.2. Acute effects of concussions

To examine the acute effects of concussion on tract microstructure, we analyzed football players who had a concussion and scanned soon thereafter during the study (n=12 players). Two main effects were tested in a longitudinal linear mixed-effects model within subjects and across nodes, 1) comparing baseline scans to scans immediately following a player’s concussion (concussion scans) (average time after concussion = 1.8± 1.3 days, range 1-5, N=11), and 2) comparing concussion scans to follow-up scans (average time from concussion to follow-up scan = 165± 77 days, range 58-280, N=10). **Figure 5** demonstrates examples of longitudinal metric changes within fiber tracts for the two effects (baseline vs. concussion and concussion vs. follow-up). Acutely after concussion, we observed a localized increase in AD in both the left uncinate and right cingulum hippocampus. This significant increase was also observed in MK and RK within the same tracts. The increase in diffusion metrics was often transient and decreased significantly at the follow-up scan (**Fig 5**a; **Suppl. Table 3**).

**Figure 5.**
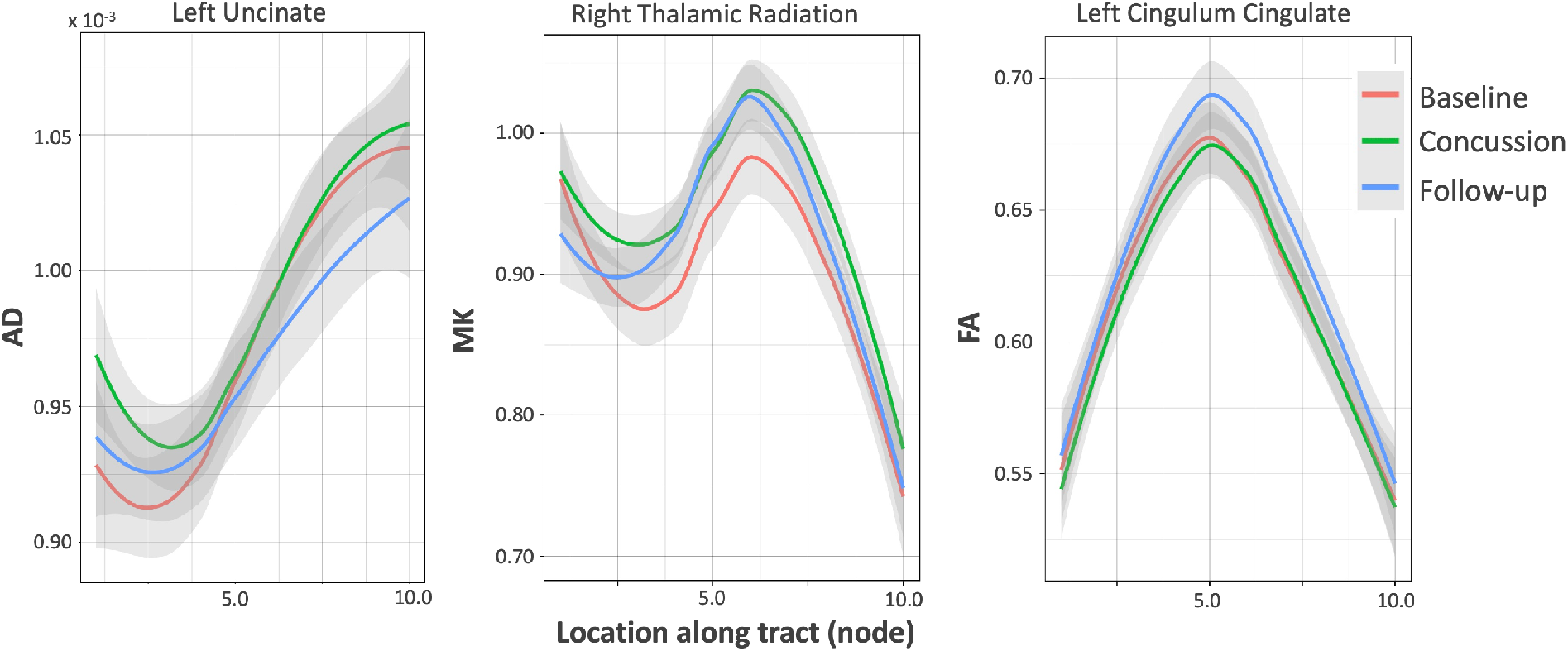
Acute effects of concussion assessed by comparing i) baseline to immediately post-concussion (concussion) scans, and ii) concussion to follow-up scans. Examples of diffusion metric changes across nodes in baseline, concussion, and follow-up scans: changes in AD within the left uncinate tract (left) represent a finding exhibiting both effects: baseline vs. concussion (T-statistic: 3.9; p-value: < 0.001) and concussion vs. follow-up (T-statistic: −4.2; p-value: < 0.001) (**Suppl. Table 3**). MK differences in the right thalamic radiation (middle) represent changes from baseline to concussion (T-statistic: 4.05; p-value: < 0.001), while FA differences in the left cingulum cingulate (right) represent changes from concussion to follow-up (T-statistic: 5.69; p-value: < 0.001).

### 3.4. Effect of player position

To further dissect the effects of sub-concussive impacts on chronic microstructural changes in high-contact sports, we investigated the association between localized microstructural changes in football players with player position-based head impact risk, as assessed by HITsp (which quantifies a player’s theoretical position-based concussion risk based on hit impact velocity, quantity, duration, and location for other players of the same position). We also studied the associations of behavioral and demographic measures (baseline SCAT, prior football exposure and the presence of prior concussion) with dMRI metrics in football players. The left cingulum hippocampus showed a group difference (main effect) where football players with high position-based sub-concussive risk had higher ODI (t=3.9, p=0.0002, **Suppl. Fig. 2**-left). The right SLF showed an interaction between time and position-based concussion risk, where players with higher concussion risk had a greater increase in ODI over time (t=3.6, p=0.0003, **Suppl. Fig. 2**-right). No other concussion risk (HITsp) effects passed multiple comparison correction. Baseline SCAT score, prior football exposure and prior concussion exposure similarly did not show a statistical relationship to dMRI metrics.

## 4. Discussion

This study examined brain microstructure in high-contact (football) and low-contact (volleyball) college athletes across multiple seasons using advanced diffusion MRI microstructural metrics and fiber quantification tools. In volleyball athletes, we found decreasing RD, MD, and ODI, and increasing FA and AWF, in multiple tracts including callosum forceps minor, left SLF, left thalamic radiation, and right cingulum hippocampus, while in football players these changes were relatively attenuated or absent. Similar effects were observed when volleyball athletes were compared to football players with a prior or in-study concussion, who showed similar relative absence of FA/AWF increases and RD/ODI decreases over time. These effects were more prominent and observed in a larger number of tracts specifically in concussed (previously or in-study) football players compared to volleyball. Football players who had a concussion during the study showed a transient localized increase in AD, MK, and RK in the left uncinate and right cingulum hippocampus. Finally, studying player position effect on microstructural changes in football revealed an increase in ODI over time in football players with high position-based sub-concussive risk. Overall, our results suggest that normative changes in diffusion metrics, which may reflect healthy brain developmental trajectories, may be significantly altered or absent in football athletes.

### Football results in divergent microstructural trajectories

In our longitudinal multi-season investigation with multicompartment modeling, we observed decreased tract-specific diffusivity and dispersion perpendicular to the primary pathway over time in low-contact sports (volleyball), whereas in high-contact athletes (football) these changes were attenuated. This finding is consistent with existing literature suggesting decreased tract-specific diffusivity, including RD^27^, MD^5^, and ODI^28^ in non- or low-contact compared to high-contact sports; particularly in the corpus callosum where we also observed decreasing MD, RD and increasing AWF. In addition, we observed increased FA longitudinally in volleyball compared to football, consistent with findings in studies including non-contact sports vs. a rugby cohort^29,30^. WMTI and kurtosis metrics can be sensitive markers of WM injury following head impacts^31–33^ as they enable the quantification of non-Gaussian water diffusion, and might be sensitive to effects such as inflammatory responses^34^; thus, employing them in addition to DTI and NODDI metrics enhances our ability to decipher underlying microstructural mechanisms.

### Altered white matter development

From a developmental perspective, myelination is thought to increase into adulthood and peak around 20-40 years of age^35,37–39^. In contrast, pruning has generally been thought of as completed by puberty/early adolescence^40–43^. While newer research suggests that pruning may continue, to a lesser degree, into early adulthood^44^, myelination is generally accepted to peak later in life. Myelination, as measured by DTI, has been generally observed as FA increases, with MD and RD decreases, during childhood and adolescence. The observed increasing AWF/FA and decreasing RD/ODI in our volleyball cohort over years likely still reflect normal development in this age range^35,36^. On the other hand, the relative attenuation of these effects seen in football could possibly reveal delayed myelination, altered axonal calibers, or depressed pruning processes leading to increased axonal/tract dispersion. This potential delayed maturation in the WM parallels our prior finding of divergent trajectories in the GM (cortical thinning) in football vs. volleyball players, likewise possibly reflecting altered normal cortical development^16^. Inclusion of additional myelin or demyelination sensitive imaging sequences, such as magnetization transfer imaging, in future work may further enable distinguishing these cellular and developmental processes.

### Tract-specific effects of long-term concussion exposure

Regarding chronic effects of concussion, players with a concussion (prior or in-study) had more extensive differences with larger divergences in multiple diffusion metrics and across several tracts (including callosum forceps minor, SLF and thalamic radiation) when compared to volleyball controls; whereas those without prior or in-study concussions had lesser differences compared to volleyball (**Table 3**). A more direct longitudinal comparison of concussive versus non-concussive football players showed that concussive football players have significantly divergent trajectories with relative decreases in FA/AWF and increases MD, RD and ODI over time compared to non-concussive football players (**Table 3**). Taken together, this suggests that non-concussive football players exhibit an intermediate imaging phenotype between concussive football and volleyball. These chronic findings are supported by several studies of WM abnormalities in concussed contact sport athletes in comparison to matched contact sport controls followed over 6 months or a single season, where concussed players exhibited: 1) decreased FA in tracts including the corpus callosum and cingulum hippocampus, associated with worsening impulse control^45^ and memory performance^46^; or 2) elevated MD and/or RD in the corpus callosum^47^, the SLF and corticospinal tract^48^. The corpus callosum is consistently reported as one of the most common sites of microstructural injury resulting from concussive impacts^49^. Finite element analyses demonstrate that it is sensitive to rotations and shear forces, leading to higher tract-oriented strains^50^, potentially due to lateral motion of the stiff falx^51^. Callosal injury is thought to affect perception, orientation, and reaction time^52^. Other tracts we found that commonly exhibit trauma-related injury include the thalamic radiation, cingulum and cortico-spinal tract^49^.

While several studies confirm our findings, some cross-sectional studies have reported changes in diffusion metrics in the opposite direction, i.e., elevated FA and lower MD and ODI in the concussion group^53–55^. This difference may be due to the large heterogeneity across these athletes and concussion populations at baseline, the heterogeneity of microstructural changes in mild traumatic brain injury (TBI) across different tracts or regions, and the complexity of comparing across distinct image analysis techniques (voxel-wise, skeletons, regions of interests, tract-based), and imaging protocols. These discrepancies further support the need for a longitudinal study design that accounts for baseline difference and more sophisticated diffusion metrics that have higher specificity to underlying intra and extracellular processes.

### Acute effects of concussion

Examining acute effects of concussion in football, players who had a concussion during the study showed a transient localized increase in AD, MK and RK in the left uncinate and right cingulum hippocampus after concussion, compared to their pre-concussion baseline scan. These effects appeared to only be transient and decreased significantly at follow-up. Previous work similarly found elevated AD acutely in the right corticospinal tract relative to controls at both 6 days and 6 months post-concussion^56^. Another study found elevated AD and RD in the genu of the corpus callosum within 2 weeks of TBI^57^. In contrast, reduced post-concussion AD has been reported over shorter time periods^58,59^, but without baseline scans and with different statistical techniques. Lancaster et al. also observed more widespread elevated AK^59^, consistent with our detected increased kurtosis metrics. The dynamics of microstructural changes pre- and post-injury are complex^5,60–62^, and depend on multiple factors such as severity of injury, duration of follow-up, scan intervals and assessed metrics. The transient increase in kurtosis metrics and AD observed in concussed players in our study may reflect hindered diffusion, possibly due to diffuse axonal injury, axonal beading, and/or an acute neuroinflammatory response and microglial activation.^24^

### Relationship to player position

We observed that players with higher position-based sub-concussive risk (HITsp) had a greater increase in ODI over time in the right SLF and an increased ODI in the left cingulum hippocampus at baseline. This effect mirrors some of our main findings of higher ODI slope over time in football compared to volleyball players, possibly reflecting delayed myelination and maturation, or reduced axonal pruning.

### Limitations

Our current study should be interpreted in light of some limitations. While we employed advanced mapping techniques to localize changes along the length of tracts (avoiding biases of averaging and mis-correspondences of voxel/skeleton approaches), this approach averages effects per node across tracts and subjects. Future work will assess subject-specific changes across the whole brain. Although we followed the players for a period of up to 4 years, we only focused on male athletes from a single institution, and the number of in-study concussions was limited, arguing for larger studies of more sports and institutions and both sexes. We used self-report of individual concussion history and cannot account for under-reported prior concussions. Similarly, participation in other high-contact sports, which was not assessed, should be formally quantified in future studies. We only had available the global SCAT score; future work can examine detailed SCAT sub-scores and other more detailed neurocognitive testing.

### Conclusion

Overall, our study used advanced diffusion MRI metrics based on signal representation (diffusion tensor and kurtosis) and multi-compartment modelling (NODDI and WMTI), to assess chronic and acute microstructural changes in athletes of collegiate high-impact sports, with low-impact sport athletes as a control population. We found callosum forceps minor and left SLF to be the most affected tracts, with microstructural changes in all our comparisons. While the control, low-contact population showed maturational changes that could signal higher myelination, such as lower RD, MD and ODI, and higher FA and AWF over time, these changes were attenuated in high-contact athletes. This study contributes to the body of literature examining the chronic effects of high-contact sports, leveraging a multi-year study duration, advanced metrics, and tract-specificity. Future longitudinal studies that include more sports will complement our knowledge about brain changes due to head impacts in sports, while combined prospective imaging and histopathology studies in animals or even humans^63,64^ might elucidate the pathological correlates of imaging changes.

## Data availability and access

All data associated with this study are available in the main text or the supplementary materials. All the imaging data can be shared upon request with a proposal and under a data transfer agreement.

## Acknowledgements

This research was conducted with funding from the Radiology Society of North America and American Society for Neuroradiology, the Center for Biological Imaging at Stanford, and General Electric Healthcare. This publication was also supported by the Pac-12 Conference’s Student-Athlete Health and Well-Being Initiative. The content of this article is solely the responsibility of the authors and does not necessarily represent the official views of the Pac-12 Conference or its members. We thank Dr. Andrew Hoffman, Dr. Tony Wyss-Coray, Steven Bartlinsky, Scott Anderson, Christopher Barrett, Lexie Ross, Eitan Gelber, and Anthony Pass for their assistance with study design, recruitment, and evaluation.

## Author contributions

MG, BDM and MZ designed the study and experiments. MG, BDM, MG, MK, NM and MZ analyzed and interpreted the data and wrote the manuscript. SS, ELD, CA, LAM, BB, DD, PG contributed to data acquisition, interpretation, and manuscript revision. JR contributed to statistical analyses and interpretation. GG, MW, DC contributed to funding acquisition, data interpretation, and manuscript revision.

