## Supplemental Material for "Microstructural alterations in tract development in college football: a longitudinal diffusion MRI study"

#### Methods

Image Acquisition

A 3T MRI scanner (GE MR 750, Milwaukee, WI, USA) and an 8-channel-receive head coil were used for scanning. We acquired:

- a 3D T1-weighted axial: 1mm isotropic, inversion recovery fast spoiled gradient echo Brain Volume Imaging (BRAVO), TR=7.9ms, TE=3.1ms, number of excitations (NEX) 1, imaging time=5.1mins.
- two whole-brain axial 2-D pulsed gradient spin echo product diffusion MRI acquisitions, with 60/30 directions at b=2500/800s/mm$^2$ and 6/3 b=0s/mm$^2$ images, TE=81ms, voxel size=1.875×1.875×2.0mm$^3$, 70 slices, imaging time=9:21/4:40mins.
- a field-map based on a multi-echo gradient echo 1mm 3D isotropic acquisition, to correct distortions.

Automated fiber quantification (AFQ)

AFQ automatically identifies the core of major fiber tracts in each subject, and samples tissue properties at 100 equally spaced nodes along the tract, considering values of neighboring voxels, weighed by their distance to the tract core. AFQ quantifies the following 20 major fiber tracts: Left/Right Thalamic Radiation, L/R Corticospinal, L/R Cingulum Cingulate, L/R Cingulum Hippocampus, Callosum Forceps Major, Callosum Forceps Minor, L/R inferior fronto-occipital fasciculus (IFOF), L/R inferior longitudinal fasciculus (ILF), L/R superior longitudinal fasciculus (SLF), L/R Uncinate and L/R Arcuate. The average tract core across the cohort is sampled to account for heterogeneity across subjects and remove outlier tracts.

At the node level, nine outliers were manually visually identified across six different tracts (Volleyball: 2 scans from 1 subject. Football: 8 scans from 6 subjects). For each of these scans, the entirety of the tract was removed from the analysis. The forceps major of the callosum was excluded for all subjects because of inaccuracies on tractography with visual inspection leading to marked variability in the metrics.

### Tables

***Suppl. Table 1. Comparison of tract-specific diffusion metrics in football vs. volleyball over time including the 95% confidence intervals around the coefficients of each of significant sport by time interaction.***

| Tract | Measure | Sport by Time T (p) | Confidence interval of interaction coefficient | Sport T (p) | Time T (p) |
|---|---|---|---|---|---|
| Callosum Forceps Minor | md | -3.68 (2.31E-4) | (-5.03E-6, -1.53E-6) | 2.09 (0.105) | 0.81 (0.253) |
| | rd | -4.14 (3.42E-5) | (-5.90E-6, -2.11E-6) | 1.31 (0.501) | -2.04 (3.05E-6) |
| | awf | 4.25 (2.09E-5) | (2.00E-3, -2.11E-6) | -1.55 (0.326) | 2.03 (2.37E-6) |
| Left SLF | fa | 4.55 (5.31E-6) | (2.11E-3, 5.29E-3) | 1.65 (0.040) | 0.62 (0.001) |
| | rd | -3.97 (7.16E-5) | (-2.26E-6, -7.66E-7) | -0.09 (0.674) | -1.81 (1.71E-5) |
| | odi | -3.93 (8.56E-5) | (-3.28E-3, -1.10E-3) | -2.06 (0.015) | 3.14 (0.163) |
| | awf | 6.08 (1.20E-9) | (2.08E-3, 4.06E-3) | 0.16 (0.449) | -0.44 (0.004) |
| Left Thalamic Radiation | rd | -3.66 (2.49E-4) | (-6.00E-6, -1.84E-6) | 1.92 (0.256) | -0.30 (0.016) |
| Right Cingulum Hippocampus | odi | -3.69 (2.25E-4) | (-6.39E-3, -1.96E-3) | 0.64 (0.880) | 0.55 (0.148) |
| | awf | 3.58 (3.41E-4) | (1.62E-3, 5.55E-3) | -1.20 (0.521) | 0.08 (0.035) |

***Suppl. Table 2. Longitudinal comparison between i) volleyball players, ii) football players with prior or in-study concussion (concussive), iii) with no history of concussion (non-concussive).*** *Only metrics with significant longitudinal differences ("Group * Time" interaction term, after FDR correction) are reported.*

| | Tract | Measure | Group by Time T (p) | Group T (p) | Time T (p) |
|---|---|---|---|---|---|
| **concussive football vs. volleyball** | Callosum Forceps Minor | FA | 4.34 (1.44E-5) | 0.14 (0.625) | 1.47 (1.36E-8) |
| | | MD | -6.96 (3.52E-12) | 2.46 (0.071) | 5.12 (0.449) |
| | | RD | -6.45 (1.09E-10) | 1.40 (0.518) | 1.24 (8.55E-5) |
| | | FICVF | 4.04 (5.39E-5) | -0.93 (0.548) | 1.64 (1.57E-8) |
| | | AWF | 6.00 (2.01E-9) | -1.22 (0.597) | -1.81 (0.005) |
| | Left Arcuate | MD | -3.81 (1.38E-4) | 2.29 (0.042) | 3.81 (0.082) |
| | Left SLF | FA | 4.43 (9.63E-6) | 2.07 (0.014) | -0.40 (0.001) |
| | | RD | -4.26 (2.05E-5) | 0.06 (0.757) | 0.41 (0.002) |
| | | ODI | -3.87 (1.09E-4) | -2.83 (0.001) | 2.71 (0.811) |
| | | AWF | 6.47 (9.51E-11) | -0.12 (0.603) | -2.79 (0.057) |
| | Left Thalamic Radiation | FA | 5.00 (5.88E-7) | -0.85 (0.932) | -1.31 (0.010) |
| | | MD | -4.57 (4.89E-6) | 3.04 (0.025) | 4.62 (0.031) |
| | | RD | -5.34 (9.18E-8) | 1.88 (0.300) | 2.90 (0.436) |
| | | ODI | -4.60 (4.16E-6) | 0.62 (0.815) | 2.62 (0.610) |
| | | AWF | 3.81 (1.41E-4) | -1.65 (0.359) | -2.60 (0.881) |
| | Right ILF | FA | 3.64 (2.71E-4) | -1.08 (0.681) | 0.60 (8.00E-5) |
| | | AWF | 3.61 (3.09E-4) | -1.63 (0.317) | -2.28 (0.917) |
| | Right SLF | FA | 4.05 (5.11E-5) | 1.40 (0.102) | -0.20 (0.001) |
| | | RD | -4.00 (6.38E-5) | -0.54 (0.428) | -2.65 (3.69E-12) |
| **non-concussive football vs. volleyball** | Left SLF | FA | 3.86 (1.14E-4) | 1.60 (0.050) | 1.19 (1.47E-5) |
| | | AWF | 4.40 (1.11E-5) | 0.60 (0.302) | 2.09 (4.86E-9) |
| | Right Cingulum Cingulate | MK | 4.30 (1.67E-5) | -0.89 (0.848) | -2.27 (0.718) |
| | Right Cingulum Hippocampus | FA | 3.74 (1.86E-4) | 0.22 (0.427) | 1.28 (1.34E-5) |
| **concussive vs. non-concussive football** | Callosum Forceps Minor | AWF | 3.64 (2.78E-4) | -0.17 (0.711) | -4.07 (0.054) |
| | Left Cingulum Cingulate | MD | 3.79 (1.52E-4) | -0.11 (0.700) | -4.16 (0.060) |
| | | RD | 4.35 (1.36E-5) | 0.41 (0.334) | -2.50 (0.092) |
| | Left SLF | FICVF | 3.59 (3.34E-4) | -0.54 (0.853) | -1.30 (0.008) |
| | Left Thalamic Radiation | FA | 4.02 (5.74E-5) | -1.86 (0.306) | -4.95 (0.004) |
| | | ODI | -3.92 (8.83E-5) | 1.54 (0.476) | 3.29 (0.811) |
| | Right Arcuate | AWF | 3.61 (3.08E-4) | -0.55 (0.836) | -5.88 (5.19E-7) |

***Suppl. Table 3. Acute effects of concussions, comparing i) baseline to immediately post-concussion (concussion) scans, and ii) concussion to follow-up scans.*** *T-statistic (p-value) for the different variables of interest from the longitudinal linear mixed-effects models for both comparisons. Metrics that exhibited significant changes for both effects within the same tract are highlighted in gray.*

| **Baseline vs. Concussion scan** | | | **Concussion vs. Follow-up scan** | | |
|---|---|---|---|---|---|
| Tract | Metric | Concussion Effect T (p) | Tract | Metric | Concussion Effect T (p) |
| Left Uncinate | AD | 3.90 (9.44E-5) | Left Uncinate | AD | -4.21 (2.50E-5) |
| | FA | 3.84 (1.25E-4) | Left Cingulum Cingulate | FA | 5.69 (1.27E-8) |
| Right Thalamic Radiation | MK | 4.05 (5.18E-5) | | RD | -5.88 (4.09E-9) |
| Right Cingulum Cingulate | FICVF | 4.17 (3.11E-5) | | ODI | -5.43 (5.77E-8) |
| Right SLF | FA | 4.19 (2.76E-5) | Left Cingulum Hippocampus | RD | -4.38 (1.17E-5) |
| | RD | -6.61 (6.47E-5) | Left Arcuate | RD | -4.03 (5.57E-5) |
| | FICVF | 5.51 (3.60E-8) | | FICVF | 3.84 (1.25E-4) |
| | AWF | 4.18 (2.94E-5) | | AWF | 5.44 (-5.33E-08) |
| Right Cingulum Hippocampus | MK | 4.13 (3.62E-5) | Right Cingulum Hippocampus | MK | -3.77 (1.64E-4) |
| | RK | 4.00 (6.47E-5) | | RK | -4.01 (6.07E-5) |

### Figures

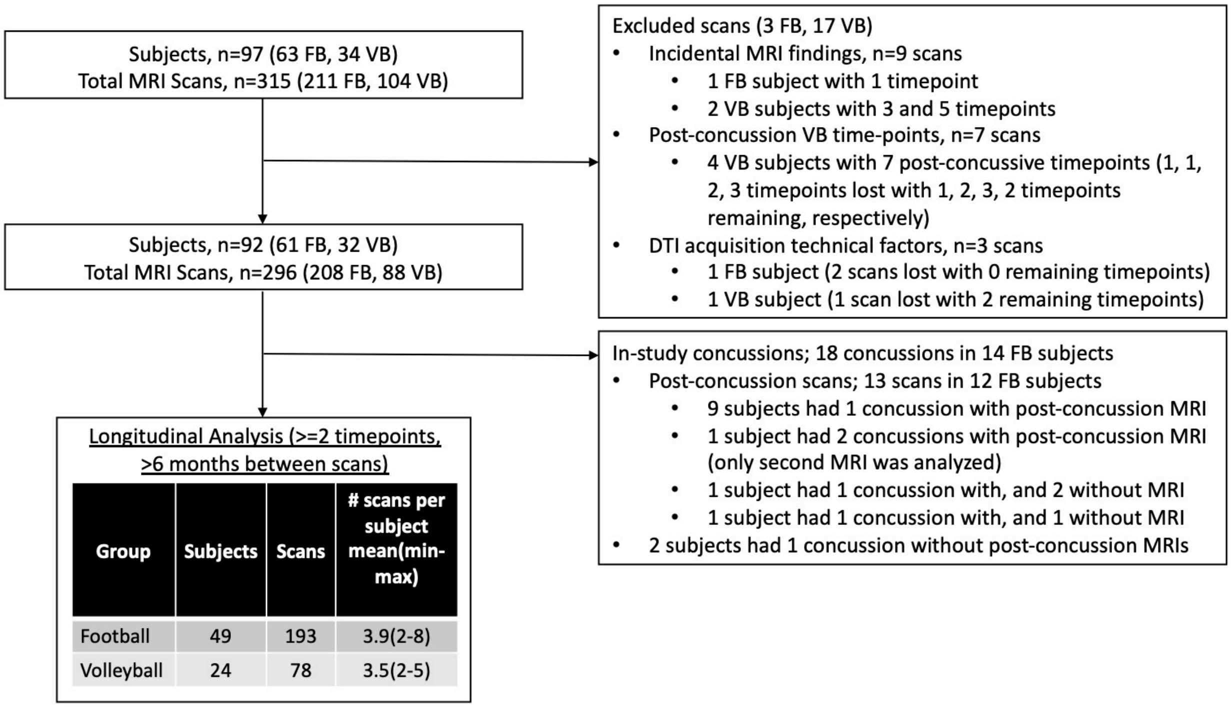


***Suppl. Figure 1**. Study enrollment showing exclusion criteria and final sample size for longitudinal analysis.* *The final analysis cohort included 49 football (FB) players and 24 volleyball (VB) players, each of which had an average of 3.9 and 3.5 scans (193 and 78 total scans in FB and VB groups, respectively).*

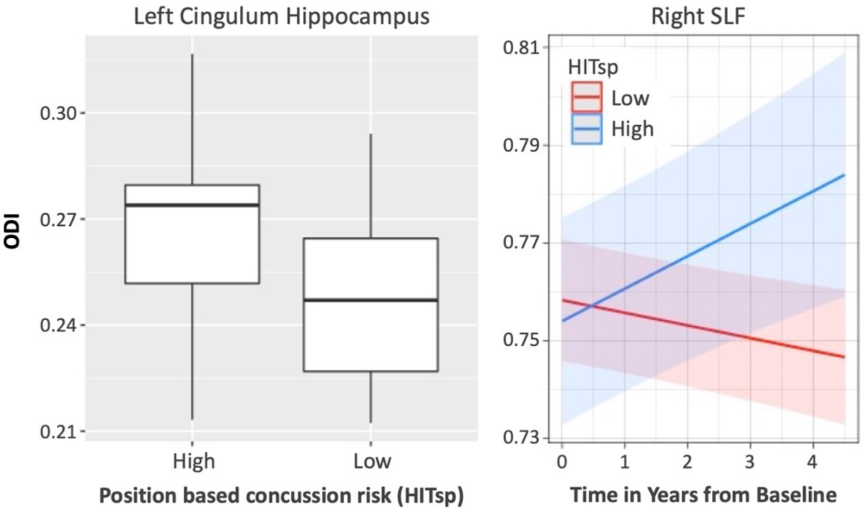


***Suppl. Figure 2*. Relationships between ODI and positional sub-concussive risk (HITsp).** <u>Left - Main effect</u>: The left cingulum hippocampus showed a group difference (main effect) where football players with high position-based sub-concussive risk had higher ODI. <u>Right – Interaction</u>: The right SLF showed an interaction between time and position-based concussion risk, where players with higher concussion risk had a greater increase in ODI over time ODI: orientation dispersion index, from NODDI. HITsp: a metric quantifying a player's position-based concussion risk based on hit impact velocity, quantity, duration, location, etc. SLF: superior longitudinal fasciculus. No other HITsp effects passed multiple comparison correction.
